# ZNHIT1-dependent H2A.Z deposition at meiotic prophase I underlies pachytene gene expression and meiotic progression during male meiosis

**DOI:** 10.1101/2024.06.06.597721

**Authors:** Shenfei Sun, Yamei Jiang, Ning Jiang, Qiaoli Zhang, Hongjie Pan, Fujing Huang, Xinna Zhang, Yuxuan Guo, Xiaoyu You, Kai Gong, Wei Wei, Hanmin Liu, Zhenju Song, Yuanlin Song, Xiaofang Tang, Miao Yu, Runsheng Li, Xinhua Lin

## Abstract

Accurate meiotic progression is important for gamete formation and the generation of genetic diversity. However, little is known about the identity of chromatin regulators that underlie mammalian meiosis in vivo. Here, we identify the multifaceted functions of the chromatin remodeler ZNHIT1 in governing meiosis. The expression of *Znhit1* gradually increases during the meiotic prophase, and *Znhit1* knockout in spermatocytes results in arrested pachytene development, impaired DNA double-strand break repair, and defective homologous recombination. Single-cell RNA sequencing and transcriptome analysis reveal that *Znhit1* loss dysregulates meiotic transcriptional programs at the pachytene stage. Chromatin immunoprecipitation data show that ZNHIT1 is needed for the incorporation of the histone variant H2A.Z into pachytene chromatin. Moreover, we found that H2A.Z cooperates with the transcription factor A-MYB to co-bind DNA elements and control gene activity. Our findings provide insights into the regulatory mechanisms governing meiotic progression and highlight ZNHIT1 as a critical regulator of meiotic progression.

## Introduction

Meiosis is a fundamental and conserved process that plays a crucial role in gamete formation and the generation of genetic diversity. By undergoing one round of DNA replication and two rounds of cell division, the meiotic process ensures the production of haploid cells, each with a unique combination of genetic materials (Handel and Schimenti, 2010; Lascarez-Lagunas et al., 2020). Any disruptions to the normal progression of meiosis would have significant consequences, such as aneuploidy, infertility, spontaneous abortion, and congenital diseases (Hassold and Hunt, 2001). Therefore, understanding the molecular mechanisms underlying meiosis will provide valuable insights for the diagnosis and treatment of reproductive and developmental diseases.

Multiple chromosomal events occur during meiotic prophase I, including homologous chromosome pairing and synapsis, DNA double-strand break (DSB) formation and repair, and homologous recombination (HR) (Baudat et al., 2013; Hunter, 2015; Zickler and Kleckner, 2023). In this process, homolog pairing and synapsis coincide with SPO11-mediated genome-wide formation of DSBs (Bergerat et al., 1997; Keeney et al., 1997). Following the formation of DSBs, the DNA undergoes resection, leading to the generation of single-stranded DNA overhangs. These overhangs are then coated by RPA, DMC1, and RAD51, facilitating the production of recombination intermediates. Subsequently, these intermediates are processed and resolved, ultimately forming either meiotic crossovers or non-crossovers (Gray and Cohen, 2016; San Filippo et al., 2008; Symington, 2014). The tightly coordinated timing and spatial arrangement of these meiotic events are of utmost importance for proper germ cell development.

There is transcriptional awakening during male meiotic prophase I, in which the meiotic genome becomes transcriptionally active during the zygotene-to-pachytene transition (Alexander et al., 2023; Ernst et al., 2019; Green et al., 2018; Rabbani et al., 2022; Turner, 2015). This process, referred to as pachytene genome activation (PGA) in this context, is responsible for the expression of numerous genes and plays an essential role in controlling meiotic and post-meiotic events. Previous studies have emphasized the significance of the transcription factors (TFs) A-MYB, BRDT, and TCFL5 in activating transcription during pachytene (Alexander et al., 2023; Bolcun-Filas et al., 2011; Cecchini et al., 2023; Gaucher et al., 2012; Li et al., 2013; Maezawa et al., 2020; Manterola et al., 2018; Ozata et al., 2020; Yu et al., 2021). Additionally, extensive chromatin remodeling takes place in spermatocytes during meiotic prophase I, involving histone variant exchange, histone modifications, and high-order genome rearrangement (Kota and Feil, 2010; Wang et al., 2017; Zheng and Xie, 2019). It has been known that SETDB1-mediated H3K9 trimethylation (H3K9me3) is required for sex chromosome transcription silencing (a process called meiotic sex chromosome inactivation, MSCI) and male meiotic procession (Hirota et al., 2018). However, the specific involvement of chromatin regulators in gene activation on autosomes remains poorly understood.

Zinc finger HIT-type containing 1 (ZNHIT1), an evolutionarily conserved subunit of the SRCAP chromatin remodeling complex, acts as a key regulator for the histone variant H2A.Z deposition to control gene expression and cell fate determination (Cai et al., 2005; Cuadrado et al., 2007; Feng et al., 2022). Recent studies have shown that ZNHIT1 is involved in a wide range of developmental processes, including muscle and lens differentiation, heart development, lung branching, and adult tissue stem cell maintenance (Cuadrado et al., 2010; Lu et al., 2022; Sun et al., 2020; Wei et al., 2022; Xu et al., 2021; Zhao et al., 2019). In our previous study, *Znhit1* deletion in early male germ cells impaired the mitosis-to-meiosis transition by regulating *Meiosin* expression (Sun et al., 2022). However, the in vivo function of ZNHIT1 in regulating meiotic progression remains unknown.

To delve into the function of ZNHIT1 in meiosis, we analyzed *Znhit1* expression and observed upregulation specifically during meiotic prophase. Here, we examine the role of ZNHIT1 in multiple meiotic events and show that *Znhit1* deletion in spermatocytes causes defects of pachytene gene expression, thereby resulting in meiotic developmental arrest. Hence, our study highlights the essential role of ZNHIT1 in ensuring meiotic progression.

## Results

### *Znhit1* expression is upregulated during meiotic prophase and *Znhit1* knockout in spermatocytes disrupts spermatogenesis

To identify potential chromatin regulators in meiosis, we first queried for chromatin factors highly expressed during the zygotene stage based on published scRNA-seq data (Fig S1A and Table S1) (Chen et al., 2018). Quantitative analysis showed that the chromatin remodeler *Znhit1* was among the most highly expressed factors (Fig 1A, B, S1B, C). RNA in situ hybridization validated that *Znhit1* expression in male germ cells decreased after meiotic initiation, followed by a gradual increase in spermatocytes from the zygotene stage to the metaphase stage (Fig 1C). These results suggest that ZNHIT1 is a potential chromatin regulator of meiotic progression.

**Figure 1.**
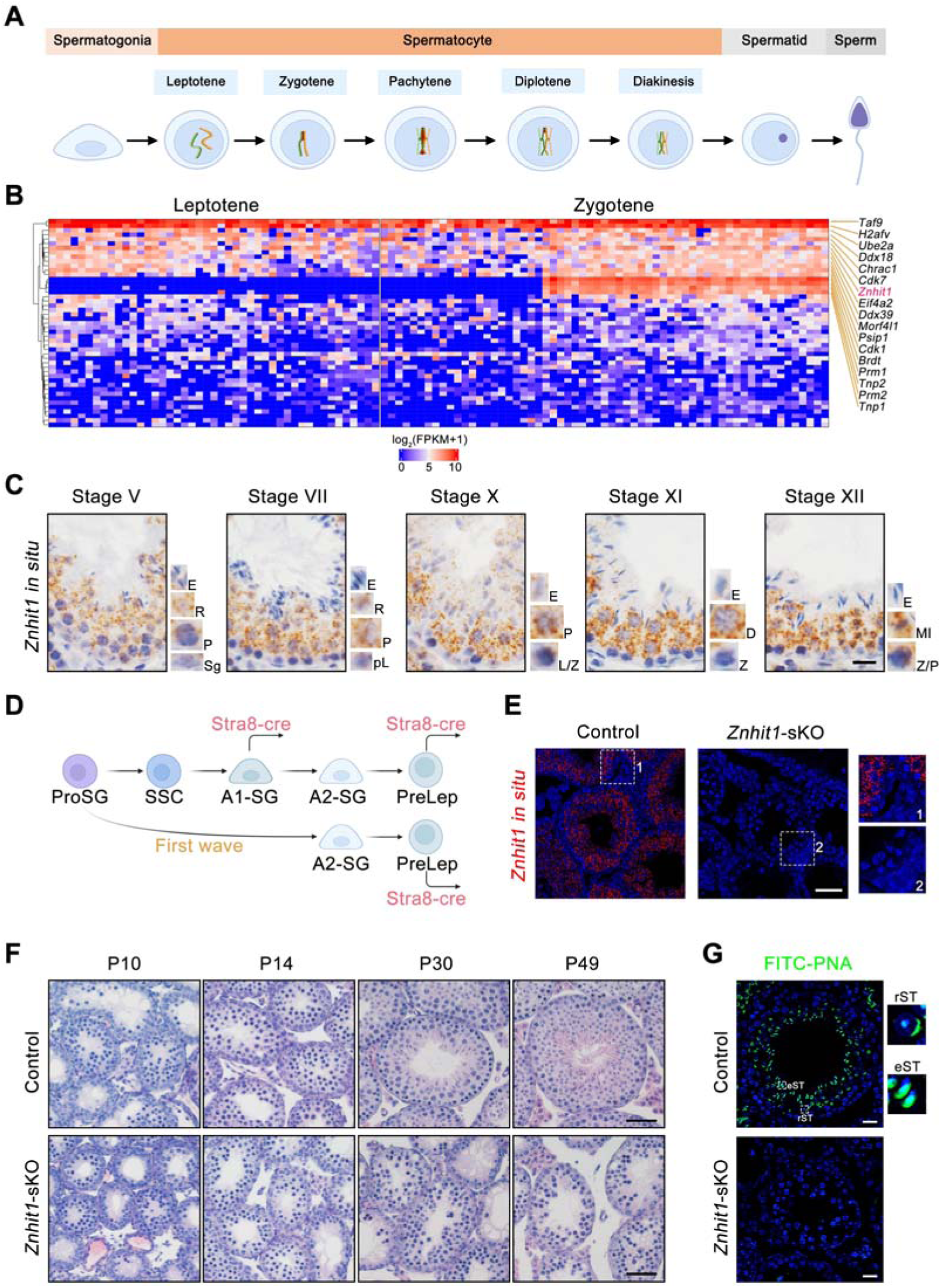
*Znhit1* deletion in spermatocytes disrupts spermatogenesis. (**A**) Schematic representation of the male germline development in mice. (**B**) Heatmap showing the clustering of 47 chromatin factors based on the transcription levels in leptotene and zygotene spermatocytes. (**C**) *Znhit1* in situ hybridization in postnatal day 60 (P60) testis sections. Scale bar, 20 μm. Sg, spermatogonia; pL, preleptotene; L, leptotene; Z, zygotene; P, pachytene; D, diplotene; MI, metaphase; R, round spermatids; E, elongating spermatids. (**D**) Schematic diagram showing the onset of Stra8-cre expression in the male germline development. (**E**) *Znhit1* in situ hybridization in P42 testis sections. Scale bar, 50 μm. (**F**) Histological testicular sections in control or *Znhit1-*sKO testis sections at the indicated times. Scale bar, 50 μm. (**G**) Immunostaining of PNA in testis sections from control or *Znhit1-*sKO mice at the indicated times. rST, round spermatids; eST, elongating spermatids. Scale bar, 20 μm. All images are representative of n = 3 mice per genotype.

To investigate whether ZNHIT1 regulates meiotic progression, we generated spermatocyte-specific *Znhit1* knockout mice (*Znhit1*^fl/fl^; *Stra8*-*cre*, referred to as *Znhit1*-sKO). In the first wave of spermatogenesis, Ngn3-negative prospermatogonia directly differentiate into A2 spermatogonia(Law et al., 2019; Rabbani et al., 2022; Yoshida et al., 2006; Yoshida et al., 2004). With Stra8-cre inducing recombination in A1 spermatogonia and preleptotene spermatocytes separately(Lin et al., 2017), it is feasible to generate mice with spermatocyte-specific gene knockout (Fig 1D). RNA in situ experiments on testis sections confirmed the deletion of *Znhit1* mRNA in spermatocytes (Fig 1E, S1D). Compared with littermate controls, *Znhit1*-sKO male mice had smaller testes (Fig S1E, F). PAS-histological and PNA-fluorescent staining illuminated the absence of round spermatids and elongated spermatids in *Znhit1*-sKO testes (Fig 1F, G), indicating that *Znhit1* knockout in spermatocytes disrupted spermatogenesis.

### *Znhit1* deletion leads to meiotic arrest at the pachytene stage

To study spermatocyte development in *Znhit1*-sKO testes, we co-stained SYCP3, a marker of primary spermatocytes, and HSPA2, a testis-specific HSP70 family member that accumulates from the pachytene stage onward. Both control and *Znhit1*-sKO testis sections exhibited SYCP3^+^HSPA2^+^ primary spermatocytes (Fig S2A). Phospho-histone H3 (Thr3) (pH3) is a marker of mitotic and meiotic metaphase cells, and *Znhit1* deletion resulted in the complete absence of pH3^+^ metaphase spermatocytes within the seminiferous tubule lumen (Fig S2B). These results suggest that *Znhit1* deletion arrests spermatocytes in meiotic prophase I.

To pinpoint the onset of meiotic failure, we performed immunostaining against SYCP3 in spermatocyte chromosome spreads. The wild-type testes displayed typical spermatocytes at consecutive substages, including leptotene, zygotene, pachytene, diplotene, and diakinesis (Fig 2A). However, spermatocytes at the diplotene and diakinesis stages were rarely observed in *Znhit1*-sKO testes (Fig 2B, C). We also examined the expression of H1T, a middle pachytene marker, and no difference in H1T expression was observed in *Znhit1*^−/−^ spermatocytes (Fig S2C). *Znhit1* knockout resulted in increased TUNEL-positive spermatocytes, suggesting that these defective spermatocytes were eliminated by the apoptotic pathway (Fig 2D, E). We further performed scRNA-seq analysis of postnatal day 35 (P35) testes from control and *Znhit1*-sKO. Uniform manifold approximation and projection (UMAP) showed that germ cells normally progressed through all meiotic stages and successfully gave rise to spermatids in control groups (Fig 2F). By contrast, in the *Znhit1* knockout group, late pachytene spermatocytes decreased significantly, and only very few subsequent germ cell types were observable. *Pou5f2* is expressed explicitly at the diplotene stage, and we found a lack of *Pou5f2*^+^ diplotene spermatocytes in *Znhit1*-sKO testes (Fig 2G). These results indicate that *Znhit1* deletion leads to meiotic arrest at the pachytene stage.

**Figure 2.**
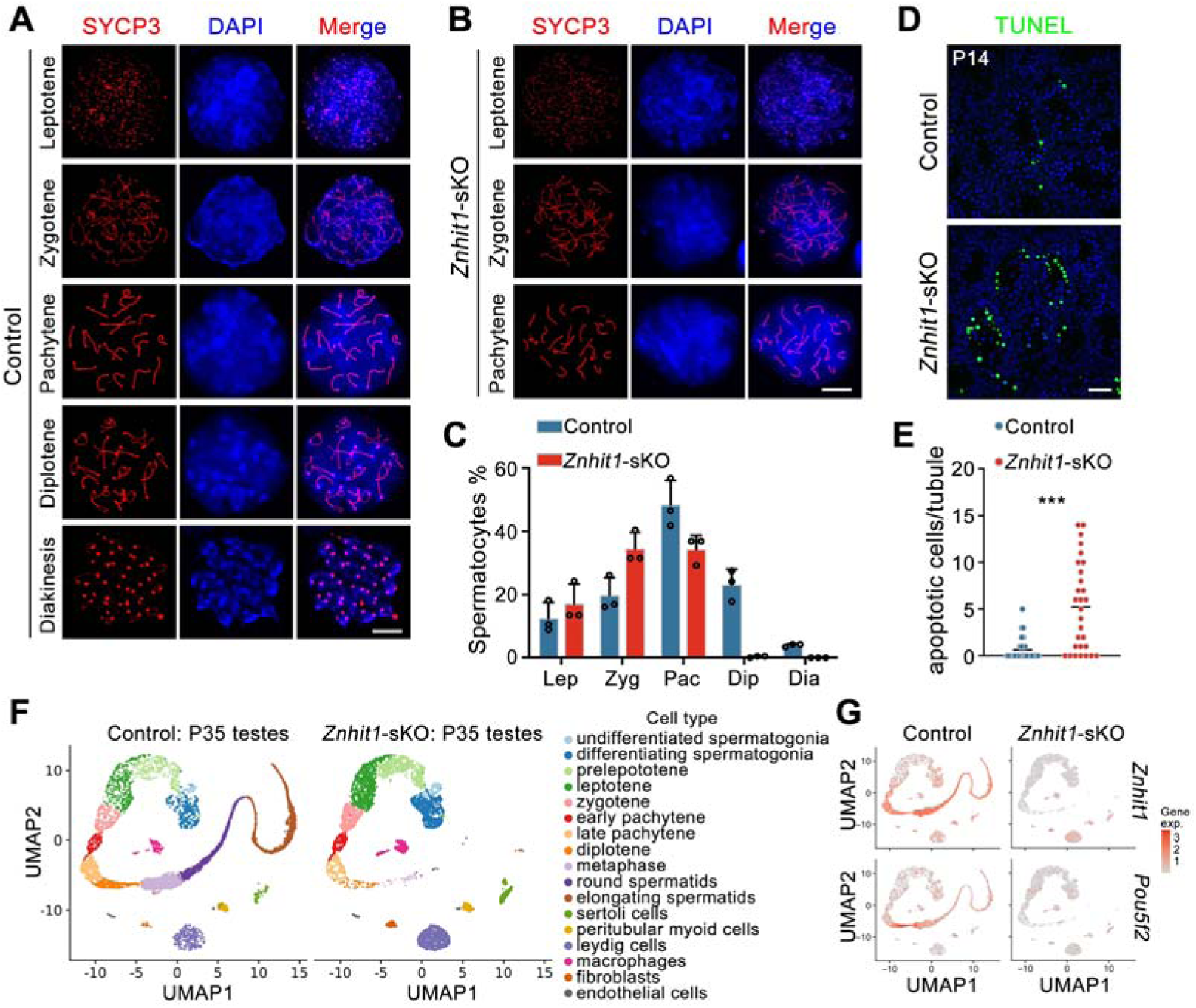
*Znhit1* deletion leads to meiotic pachytene arrest. (**A**-**C**) Immunostaining of SYCP3 in spermatocyte chromosome spreads of control (**A**) or *Znhit1*-sKO mice (**B**). Quantitative data are shown in (**C**). Scale bar, 10 μm. (**D** and **E**) TUNEL staining in testis sections from P14 control or *Znhit1*-sKO mice. Quantitative data are shown in (**E**). Scale bar, 40 μm. All images are representative of n = 3 mice per genotype. (**F**) UMAP plot showing the annotated cells captured from P35 control and *Znhit1*-sKO testicular cells. (**G**) UMAP plot showing the expression of *Znhit1* and *Pou5f2* in control P35 control and *Znhit1*-sKO testicular cells. Data are presented as the mean ± s.d. *** *p* < 0.001.

### ZNHIT1 is required for DSB repair and meiotic recombination

To investigate the impact of ZNHIT1 on meiotic events, we first performed immunostaining of SYCP1 and SYCP3, the essential components of the synaptonemal complex, to examine whether ZNHIT1 regulates homologous synapses. Pachytene nuclei with fully synapsed autosomes were observed in spermatocytes from the control (Fig. 3A). In *Znhit1*-sKO testes, while autosomal synapsis appeared grossly normal, abnormalities were present at the terminal ends of autosomes. During the pachytene stage, the X and Y chromosomes undergo pairing and synapses between the short pseudoautosomal regions (PARs). We found an increase in the percentage of unsynapsed X-Y chromosomes in *Znhit1*-sKO spermatocytes (Fig 3B). Thus, deletion of *Znhit1* results in impaired chromosomal synapsis.

**Figure 3.**
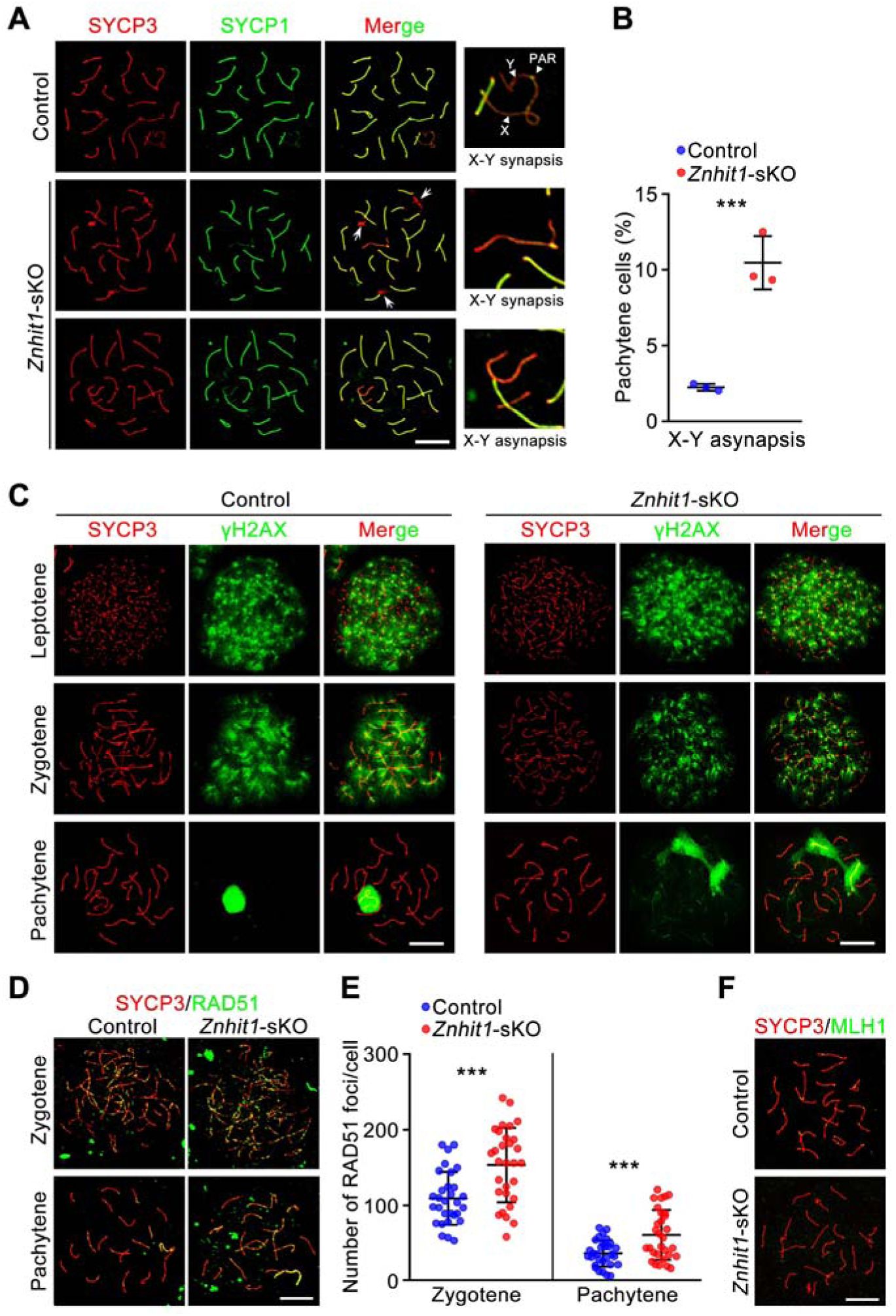
*Znhit1* deletion impairs meiotic recombination. (**A** and **B**) Immunostaining of SYCP3 and SYCP1 in spermatocyte chromosome spreads of control or *Znhit1*-sKO mice. Arrows indicate autosomal regions with abnormal synapsis. X, X chromosome; Y, Y chromosome; PAR, pseudoautosomal region. Quantitative data are shown in (**B**). Scale bar, 10 μm. (**C**) Immunostaining of SYCP3 and γH2AX in spermatocyte chromosome spreads of control or *Znhit1*-sKO mice. Scale bar, 10 μm. (**D** and **E**) Immunostaining of SYCP3 and RAD51 in spermatocyte chromosome spreads of control or *Znhit1*-sKO mice. Quantitative data are shown in (**E**). Scale bar, 10 μm. (**F**) Immunostaining of SYCP3 and MLH1 in spermatocyte chromosome spreads of control or *Znhit1*-sKO mice. Scale bar, 10 μm. All images are representative of n = 3 mice per genotype. Data are presented as the mean ± s.d. *** *p* < 0.001.

Next, we examined the programmed formation and repair of DSBs by co-staining SYCP3 and γH2AX, a DSB marker, on spermatocyte chromosome spreads. In both control and *Znhit1*-sKO spermatocytes, γH2AX signals were evident in the leptotene and zygotene stages (Fig 3C). However, in *Znhit1*-deficient pachynema, we still observed diffuse γH2AX signals on the autosomes, while in control pachynema, γH2AX signals only accumulated on the X-Y chromosomes. These results demonstrate that ZNHIT1 is required for completing DSB repair.

To understand how ZNHIT1 regulates DSB repair, we performed an immunostaining analysis using the markers RPA2 and RAD51, the essential factors involved in DSB repair. We found an increase in RPA2 and RAD51 counts in both zygonema and pachynema of *Znhit1*-deficient testes (Fig 3D, E, and S3A, B), indicating that *Znhit1* deletion delayed recombinational repair, resulting in defective resolution of autosomal DSBs in *Znhit1*^−/−^ pachynema. The formation and repair of DSBs play crucial roles in initiating and facilitating meiotic recombination, respectively. To examine the consequences of unrepaired DSBs on meiotic recombination in *Znhit1* mutants, we performed an immunostaining analysis against MLH1, a marker of meiotic crossover formation. Indeed, we found an almost absence of MLH1 foci in *Znhit1*^−/−^ pachytene spermatocytes compared with controls (Fig 3F and S3C), indicating that *Znhit1* deletion severely disrupted meiotic crossover formation.

### scRNA-seq analysis reveals that *Znhit1* deletion impairs meiotic transcriptional programs

We next investigated whether *Znhit1* deletion impacts meiotic gene expression programs. To address this, we isolated testicular cells from control and *Znhit1*-sKO mice at P16 and performed scRNA-seq analysis. scRNA-seq analysis identified 6 distinct germ cell types based on the expression of canonical stage-specific markers (Chen et al., 2018), including undifferentiated spermatogonia (*Zbtb16*), differentiating spermatogonia (*Kit*), preleptotene spermatocytes (*Nacad*), leptotene spermatocytes (*Meiob*), zygotene spermatocytes (*Psma8*), and pachytene spermatocytes (*Piwil1*) (Fig 4A and S4A, B). We then analyzed the average expression of gene sets dynamically changed during meiotic transitions, including genes upregulated or downregulated at the leptotene-to-preleptotene (Lep/PreL), zygotene-to-leptotene (Zyg/Lep), and pachytene-to-zygotene (Pac/Zyg) stages (Table S2). While *Znhit1* deletion had minimal effects on genes activated or repressed during early meiotic transitions (PreL→Lep and Lep→Zyg), it specifically and severely impaired the transcriptional programs at the zygotene-to-pachytene transition. Specifically, the average expression of 1,560 genes was activated at the pachytene stage (referred to as pachytene-activated genes, or PGA genes), while *Znhit1* deletion downregulated these PGA genes (Fig 4B). Functional enrichment analysis revealed that these PGA genes were primarily involved in DNA repair, cilium organization, and spermatid development, consistent with the biological process of germ cell development (Table S3).

**Figure 4.**
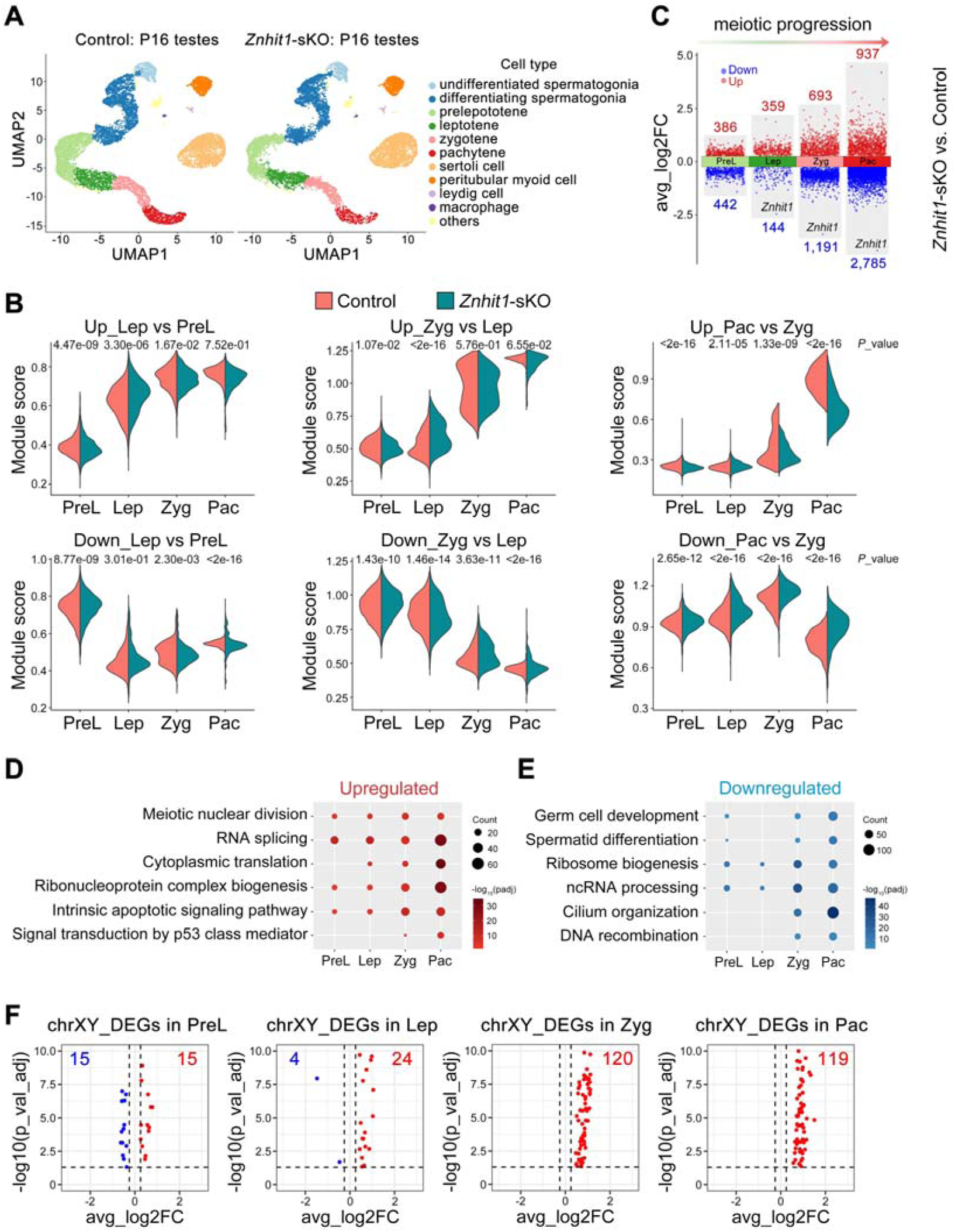
*Znhit1* deletion impairs meiotic transcriptional programs. (**A**) UMAP plot showing the annotated cells captured from P16 control and *Znhit1*-sKO testicular cells. (**B**) Violin plots showing the average expression level of upregulated or downregulated genes between two consecutive spermatocyte stages. PreL, preleptotene; Lep, leptotene; Zyg, zygotene; Pac, pachytene. (**C**) Volcano plot showing the distribution of upregulated and downregulated DEGs for each cell type in spermatocytes between control and *Znhit1*-sKO testicular cells. (**D** and **E**) Representative shared Gene Ontology (GO) terms of upregulated and downregulated DEGs at consecutive spermatocyte stages. (**F**) Volcano plot showing differentially expressed XY-linked genes in scRNA-seq data following *Znhit1* deletion.

Moreover, we identified differentially expressed genes (DEGs) in each spermatocyte type between the control and *Znhit1*-sKO groups. The number of DEGs gradually increased from the preleptotene to the pachytene stage, with downregulated genes predominating (Fig 4C and Table S4). In particular, *Znhit1*^−/−^ pachytene spermatocytes exhibited 2,785 downregulated genes, among which were most of the pachytene-activated genes (1,094 out of 1,560, 70.1%). Functional enrichment analysis revealed that upregulated genes following *Znhit1* deletion were associated with apoptotic pathways, supporting an observation that *Znhit1* deletion resulted in germ cell clearance through apoptosis (Fig 4D). Moreover, downregulated GO terms included germ cell development and DNA recombination (Fig 4E and Table S5). Interestingly, recent studies showed that cilium organization was essential for spermatocyte development (Mytlis et al., 2022), and *Znhit1* deletion reduced the expression of cilium-related genes in zygotene and pachytene spermatocytes (Fig 4E). We further conducted bulk RNA-seq experiments with P14 control and *Znhit1*-sKO testicular cells and identified 882 DEGs (812 downregulated genes and 70 upregulated genes) in *Znhit1*-sKO testicular cells (Fig S5A, and Table S6). Through integrative analysis of bulk and single-cell RNA-seq data, we found that the expression of these 812 genes was activated during the zygotene-to-pachytene transition, and *Znhit1* deletion significantly reduced the expression of these genes (Fig S5B). We also analyzed the expression of XY-linked genes. As shown in Figure 4F, while only modest numbers of XY-DEGs were detected at preleptotene and leptotene, a striking accumulation of upregulated XY-DEGs was observed at zygotene (120 genes) and pachytene (119 genes). This aberrant activation of XY-linked genes directly reveals a failure of Meiotic Sex Chromosome Inactivation (MSCI) in *Znhit1*-knockout spermatocytes. Together, these data indicate that ZNHIT1 is essential for the regulation of meiotic transcriptional programs.

Consistent with meiotic recombination defects in *Znhit1* mutants, genes associated with homologous recombination were repressed following *Znhit1* deletion, including *Ccnb1ip1*, *Rnf212*, *Spo16*, *Ankrd31*, and *Terb1* (Boekhout et al., 2019; Papanikos et al., 2019; Qiao et al., 2014; Rao et al., 2017; Reynolds et al., 2013; Shibuya et al., 2014; Wang et al., 2019; Zhang et al., 2019) (Fig S5C, D).

### *Znhit1* deletion impairs H2A.Z incorporation into pachytene chromatin

We then asked how ZNHIT1 regulates transcription. ZNHIT1 is a subunit of the SRCAP complex that facilitates the incorporation of the histone variant H2A.Z by replacing H2A (Fig S6A)(Yu et al., 2024). Immunostaining against H2A.Z showed a gradual increase in H2A.Z accumulation on autosomes from the leptotene to the diakinesis stage, while the H2A.Z signal was markedly low on the X-Y body (Fig 5A). *Znhit1* loss decreased H2A.Z staining on the pachytene cells, suggesting that ZNHIT1 is required for maintaining H2A.Z integrity during meiosis.

**Figure 5.**
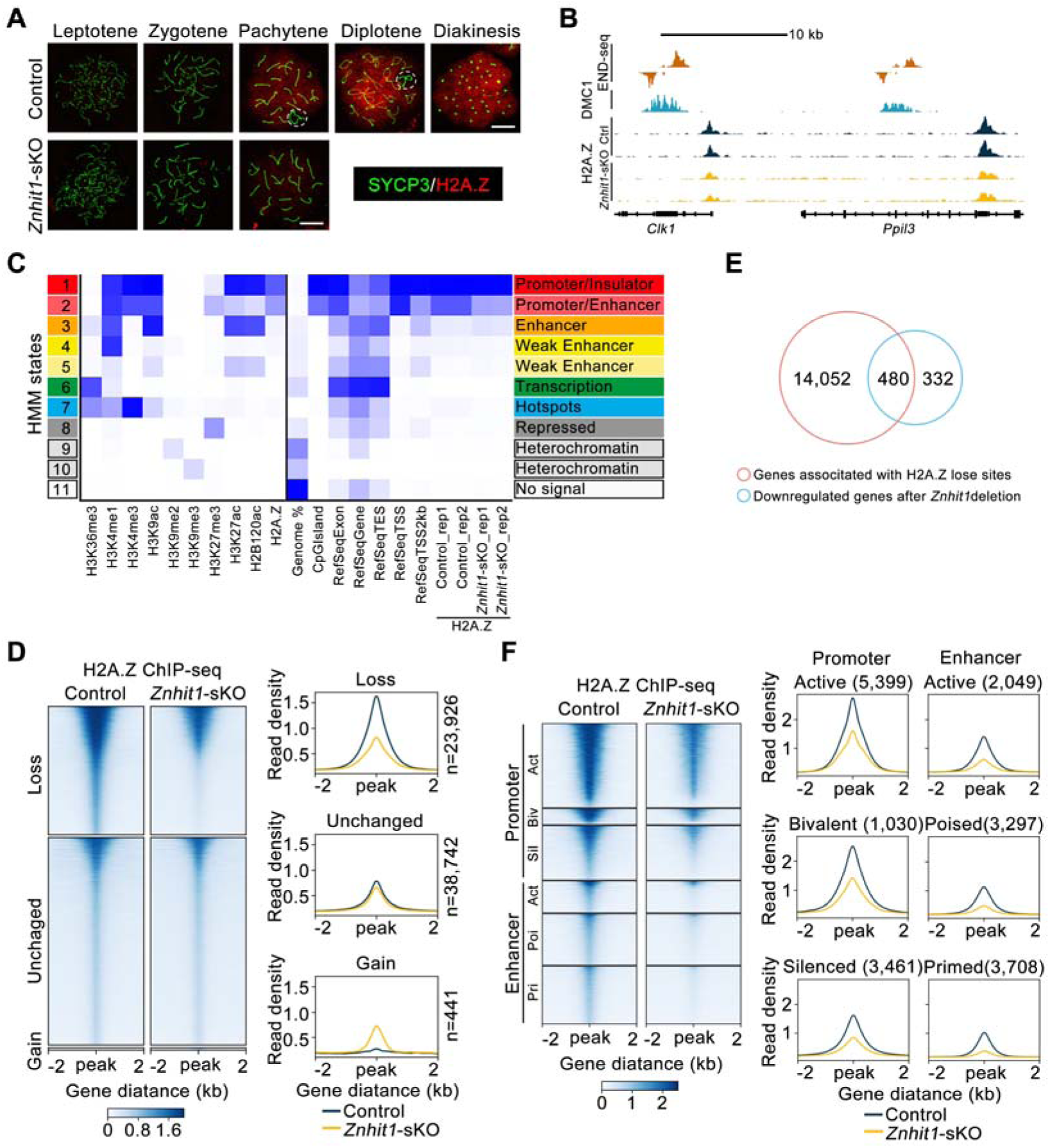
ZNHIT1 regulates H2A.Z binding on promoter and enhancer regions. (**A**) Immunostaining of SYCP3 and H2A.Z in spermatocyte chromosome spreads of control or *Znhit1*-sKO mice. The dashed circle indicates X-Y chromosomes. Scale bar, 10 μm. (**B**) Representative tracks of END-seq, DMC1-SSDS, and H2A.Z ChIP-seq in wild-type, control, or *Znhit1*-sKO testicular cells. (**C**) Heatmaps of chromatin states produced by ChromHMM and showing different enrichment for H2A.Z in control or *Znhit1*-sKO testicular cells. (**D**) Heatmaps and mean plots of H2A.Z ChIP-seq signal in control or *Znhit1*-sKO testicular cells (n = 2). (**E**) Venn diagram showing the overlap between decreased H2A.Z-bound genes and downregulated genes in *Znhit1*-sKO testicular cells. (**F**) Heatmaps and mean plots of downregulated H2A.Z ChIP-seq signals surrounding different types of promoters and enhancers.

To profile H2A.Z target genes, we performed chromatin immunoprecipitation followed by sequencing (ChIP-seq) against H2A.Z in P14 testicular cells. Using the ChromHMM model to annotate H2A.Z signals with meiocyte chromatin states defined by published epigenetic markers (Spruce et al., 2020), we found that H2A.Z sites were primarily enriched in chromatin state 1 and state 2 (promoters and enhancers) but were less enriched in recombination hotspots (Fig 5B, C), consistent with previously published results (Spruce et al., 2020). Quantitative analysis showed significant H2A.Z reduction at 37.9% (23,926 of 63,109) of H2A.Z-bound genome sites following *Znhit1* deletion (Fig 5D). Comparison analysis with bulk RNA-seq data identified 480 downregulated DEGs with decreased H2A.Z signals (Fig 5E). ∼41.3 and 37.8% of the downregulated H2A.Z peaks occupied promoters and enhancers, respectively. At promoters, these downregulated H2A.Z preferentially occupied active promoters (H3K4me3 only), while H2A.Z occupied three types of enhancers: active, poised, and primed enhancers (Fig 5F, and S6B, C). These findings indicate that *Znhit1* deletion impairs H2A.Z incorporation into pachytene chromatin.

### H2A.Z deposition cooperates with A-MYB to regulate transcription

We further asked how ZNHIT1/H2A.Z deposition facilitates lineage-specific gene expression. We performed TF motif analysis using active promoters/enhancers with decreased H2A.Z signals. This analysis identified several MYB family TFs, including MYB, A-MYB, and B-MYB (Fig 6A). Gene regulatory network analysis conducted by SCENIC further identified A-MYB (encoded by the *Mybl1* gene) as the core regulatory TF in spermatocytes (Fig 6B). We reanalyzed RNA-seq data in P14 testes from A-MYB-deficient mice (Li et al., 2013) and found that A-MYB-deficient DEGs correlated with those observed in *Znhit1*-deficient testicular cells (Fig 6C). Although *Znhit1* deletion didn’t affect *Mybl1* mRNA expression, transcriptome analyses showed that the expression of A-MYB target genes was significantly reduced in *Znhit1*^−/−^ pachytene spermatocytes (Fig 6D, E, and S7A). These results suggest that *Znhit1*^−/−^ and *Mybl1*^−/−^ pachytene spermatocytes share common gene expression alterations.

**Figure 6.**
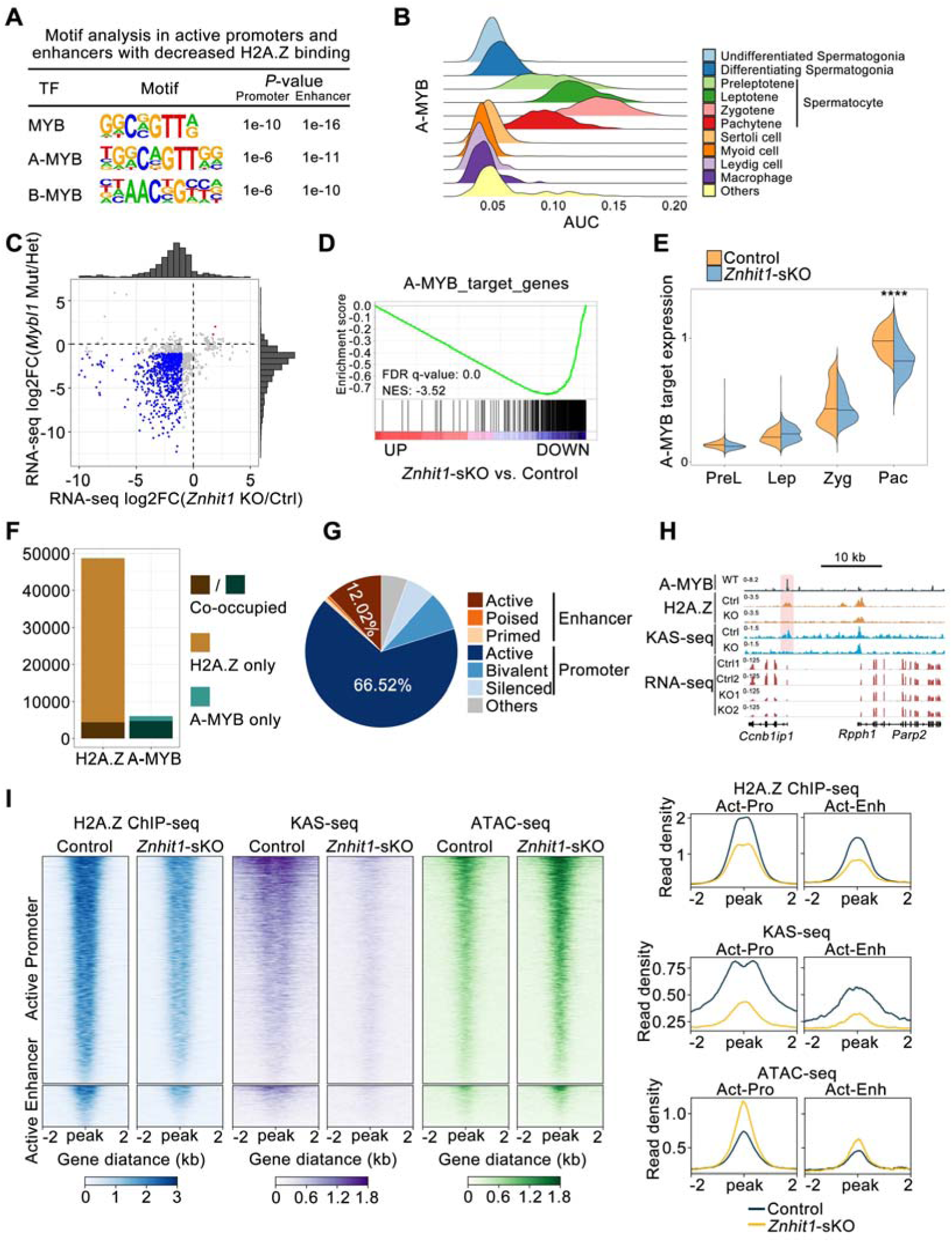
ZNHIT1/H2A.Z/A-MYB axis regulates chromatin state and gene expression. (**A**) Motif analysis of active promoters or active enhancers with decreased H2A.Z binding for putative transcription factor (TF)-binding sites using the HOMER database. (**B**) Ridge map showing the A-MYB activity (AUC) at each cell type. (**C**) Scatter plot with marginal histograms comparing the fold change of DEGs from *Mybl1*-related RNA-seq data with *Znhit1*-related RNA-seq data in P14 testicular cells. (**D**) GSEA of RNA-seq data for the control and *Znhit1*-sKO testicular cells. Selected gene sets encoded products related to A-MYB target genes. (**E**) Violin plots showing the expression of A-MYB target genes at consecutive spermatocyte stages. (**F**) Histogram showing H2A.Z and A-MYB binding sites. (**G**) Pie chart showing the distribution of H2A.Z and A-MYB co-binding sites. (**H**) Representative H2A.Z ChIP-seq tracks, KAS-seq signals, and RNA-seq signals of the *Ccnb1ip1* locus in the control and *Znhit1-*sKO testicular cells, compared with A-MYB ChIP-seq peaks. (**I**) Heatmaps and mean plots of H2A.Z ChIP-seq, KAS-seq, and ATAC-seq showing signal changes within A-MYB and H2A.Z co-binding active promoters or active enhancers. **\*\*\*\*** *p* < 0.0001.

Next, we compared the genome binding sites of H2A.Z and A-MYB. Utilizing published A-MYB ChIP-seq data from P14 testicular cells (Li et al., 2013), we identified 6,088 A-MYB binding signals, with 78.1% (4755) of these coinciding with H2A.Z peaks (Fig 6F). Annotation of genomic features showed that approximately 66.52% and 12.02% of these overlapping sites occupied active promoters and enhancers, respectively, representing a stronger enrichment compared to other regulatory elements (Fig 6G). Genes associated with H2A.Z and A-MYB binding included essential HR-related genes, such as *Ccnb1ip1* (Fig 6H). Moreover, we showed that *Znhit1* deletion resulted in a marked reduction in H2A.Z signals and nascent transcription identified by KAS-seq signals (Fig 6I). We further linked KASDseq signals with gene expression profiles, and found that *Znhit1* depletion caused a global reduction in KASDseq signals, especially at promoters of downregulated genes (Fig S8A), indicating defective transcription bubble formation. In comparison, genes with increased expression showed low KASDseq signals in both control and mutant groups, likely reflecting indirect regulation. These results further support the essential role of ZNHIT1 in transcriptional regulation.

Previous studies have shown the fundamental role of A-MYB in activating meiotic enhancers (Maezawa et al., 2020). We found that *Znhit1* deletion resulted in reduced KAS-seq signals at active enhancers (Fig 6I). Interestingly, the chromatin accessibility detected by ATAC-seq at active enhancers increased after *Znhit1* deletion, suggesting that *Znhit1* deletion results in dysregulated chromatin state. These findings suggest that ZNHIT1/H2A.Z deposition is essential for enhancer activation.

Together, these results demonstrate the critical role of the ZNHIT1/H2A.Z/A-MYB axis in governing meiotic transcriptional programs.

## Discussion

Meiosis plays a crucial role in the production of haploid gametes. Any disturbances during meiotic progress are known to lead to infertility and congenital diseases. Here we delineate the multifaceted functions of the chromatin remodeler ZNHIT1 in regulating meiotic progression. Our study provides convincing data to show that the ZNHIT1/H2A.Z/A-MYB axis controls spermatocyte development, meiotic recombination, and PGA. Thus, these findings underscore the significance of specific chromatin structures in governing appropriate transcriptional reprogramming and precise meiotic progression.

The role of H2A.Z in DNA injury and repair has been extensively studied (Colino-Sanguino et al., 2022; Dong et al., 2014; Xu et al., 2012; Yamada et al., 2018), but its function in programmed DSB formation and repair during mammalian meiosis remains unknown. To address this question, we generated *Znhit1* conditional knockout mice specifically during male meiosis to examine the role of ZNHIT1-mediated H2A.Z deposition in the formation and repair of programmed DSB. One new observation in this study is that *Znhit1* loss is dispensable for DSB formation but necessary for DNA break repair during meiosis. Intriguingly, unlike the observations in plant meiosis but consistent with Christopher L. Baker’s group observation in mouse, we confirmed that H2A.Z does not directly bind DNA recombination sites in mammalian spermatocytes (Choi et al., 2013; Spruce et al., 2020; Wang and Copenhaver, 2018). Moreover, we showed that *Znhit1* deletion delays recombinational repair but has limited impacts on DNA resection, ultimately causing defective meiotic crossover formation. Importantly, our study demonstrates that ZNHIT1 is essential for the expression of meiotic recombination-related genes, such as *Ccnb1ip1* and *Rnf212*, implying that ZNHIT1/H2A.Z modulates meiotic recombination through transcriptional regulation, rather than a direct regulatory effect of ZNHIT1 on recombination machinery. Therefore, our data reveal a different mechanism underlying meiotic recombination between plants and mammals.

Pachytene spermatocytes utilize a highly specific mechanism of meiotic surveillance, the pachytene checkpoint, to prevent aneuploidy formation by removing abnormal germ cells with incomplete chromosome synapsis and defective homologous recombination (Huang and Roig, 2023; Roeder and Bailis, 2000; Subramanian and Hochwagen, 2014). Two checkpoint pathways exist in male meiosis: one responding to unrepaired DSBs and the other triggered by MSCI failure (Royo et al., 2013). However, whether epigenetic regulation is involved in meiotic surveillance is not well defined. Previous studies have reported that upon stimulation by unrepaired DSBs, the MRE11-ATM-CHK2 pathway is activated to eliminate aberrant germ cells (Marcet-Ortega et al., 2017). Our study found that *Znhit1* deletion impairs pachytene development and XY-linked gene repression, ultimately causing meiotic pachytene arrest and apoptosis. Therefore, these findings demonstrate that ZNHIT1 acts as an important chromatin factor involved in the regulation of the pachytene checkpoint.

It has long been noticed that in meiotic prophase I, chromatin that has not yet completed chromosomal synapses is transcriptionally inactive, known as meiotic silencing of unsynapsed chromatin (MSUC) (Turner et al., 2005). When germ cells enter the pachytene stage, a large number of protein-coding genes and non-coding RNAs begin to be actively expressed, known as PGA. Previous studies have shown that transcription factors A-MYB and BRDT are involved in transcriptional activation during meiotic prophase, but how PGA takes place in the chromatin context is unclear. In this study, we found that the expression of the chromatin remodeler Znhit1 is specifically upregulated during the zygotene-to-pachytene transition. We also observed that H2A.Z deposition is enriched in the autosomes but less in the sex chromosomes at the pachytene stage. *Znhit1* deletion in spermatocytes repressed pachytene gene activation globally and reduced chromosome-wide H2A.Z deposition, supporting a central role of ZNHIT1 in PGA regulation and chromatin remodeling. It has been known that A-MYB is a transcription factor that controls pachytene transcription activation. We found that *Znhit1*-deficient spermatocytes phenocopied abnormal meiotic phenotypes observed in A-MYB mutants, such as X-Y synapsis failure, impaired DSB repair, and defective HR (Alexander et al., 2023; Bolcun-Filas et al., 2011; Li et al., 2013; Maezawa et al., 2020). Our results also revealed that ZNHIT1/H2A.Z cooperates with A-MYB to regulate PGA gene activation. Therefore, our study illuminates the molecular mechanisms underlying the fundamental question of how PGA is regulated.

The histone variant H2A.Z is enriched at gene promoters and regulatory regions, yet there is ongoing debate about its role in transcriptional regulation. A recent paper reported that H2A.Z knockout in post-mitotic muscle cells has limited effects on gene expression (Belotti et al., 2020). In this study, we utilized the meiotic prophase as a model system (where DNA replication does not occur) to study H2A.Z’s function in transcriptional regulation. We found that ZNHIT1-mediated H2A.Z deposition, independent of DNA replication, is indispensable for the transcriptional activation of a large number of meiotic genes. One explanation of this conflicting phenomenon is that H2A.Z dynamics, but not stable H2A.Z accumulation is essential for priming transcriptional changes.

Through the integration of functional and molecular evidence, our findings establish the critical involvement of ZNHIT1-dependent chromatin remodeling in the orchestration of meiotic progression and coordination of various meiotic processes, such as HR and PGA. Furthermore, we pinpoint the pivotal role played by the ZNHIT1/H2A.Z/A-MYB axis in driving transcriptional reprogramming during meiotic prophase. Taken together, this study deepens our understanding of the interplay between epigenetic regulation and mammalian meiosis.

## Materials and Methods

### Animals

*Znhit1*^fl/fl^ mice have been previously described (Zhao et al., 2019) and are available from the Model Animal Research Center of Nanjing University (MARC, Nanjing, China). The *Stra8*-*cre* knock-in mouse line was kindly provided by Dr. Ming-Han Tong(Lin et al., 2017). All mice were maintained on the C57BL/6J background. Germ cell-specific *Znhit1* knockout mice (*Znhit1*^fl/fl^; *Stra8*-*cre*) were obtained by crossing *Znhit1*^fl/+^; *Stra8*-*cre* mice with *Znhit1*^fl/fl^ mice. All mice were housed in the SPF (Specific-Pathogen-Free) animal facility with standard 12 h light/dark cycles and standard temperature (22 to 24D). All mice were provided with ad libitum access to standard laboratory food and water. All experiments in this study were performed following relevant guidelines and approved by the Animal Care and Use Committee of Fudan University.

### Histological and immunohistochemical analysis

Testes were fixed in modified Davidson’s fixative as previously described (Latendresse et al., 2002), embedded in paraffin, and sectioned. For periodic acid-Schiff (PAS)-hematoxylin staining, 5 μm testis sections were deparaffinized, rehydrated, and stained with Schiff’s reagent and hematoxylin solution. For immunofluorescent staining, 5 μm testis sections were retrieved by sodium citrate antigen retrieval buffer (pH 6.0) and blocked with 5% BSA in PBS for 30 min at room temperature. The sections were later incubated overnight at 4°C with primary antibodies as follows: mouse anti-SYCP3 (Abcam, ab97672), rabbit anti-HSPA2 (Abcam, ab108416), rabbit anti-pH3 (Millipore, H0412), rabbit anti-H1T (Invitrogen, PA5-51200), or lectin PNA (SigmaDAldrich, L7381). On the following day, secondary fluorescein-conjugated antibodies and DAPI (SigmaDAldrich, D9542) were added for 1 h, followed by Fluoromount-G mounting (Southern Biotech, 0100-01). Images were analyzed using the confocal microscope.

TUNEL staining was carried out using the DeadEnd^TM^ Fluorometric TUNEL System (Promega, G3250) according to the manufacturer’s instructions.

### Immunostaining of spermatocyte chromosome spreads

Spermatocyte chromosome spreads were prepared as previously described (Alavattam et al., 2018). Briefly, the clumps of seminiferous tubules were transferred to the hypotonic extraction buffer (30 mM Tris base, 17 mM trisodium citrate, 5 mM EDTA, 50 mM sucrose, 5 mM dithiothreitol (DTT), and 1× protease inhibitor, pH 8.2) and incubated on ice for 1.5 h. The clumps of tubules were then transferred to 30 μL of ice-cold 100 mM sucrose and mashed gently using tweezers to obtain a cell suspension. An additional 30 μL of ice-cold 100 mM sucrose was added to the cell suspension and mixed several times. Positively charged slides were incubated in the ice-cold fixation solution (2% paraformaldehyde, 0.1% Triton X-100, and 0.02% SDS, pH 9.2) for 3 min. 30 μL of the diluted cell suspension was applied to the slides and incubated in humid chambers at room temperature for 2 h. The slides were washed with 0.4% Photo-Flo 200 and stored at −80D.

For immunostaining of spermatocyte chromosome spreads, slides were washed with PBS and blocked with 5% BSA in PBS for 30 min at room temperature. The slides were stained with primary antibodies as follows: rabbit anti-SYCP3 (Abcam, ab15093), mouse anti-SYCP3 (Abcam, ab97672), rabbit anti-SYCP1 (Abcam, ab15090), mouse anti-γH2AX (BioLegend, 613401), rabbit anti-RAD51 (Proteintech, 14961-1-AP), rabbit anti-RPA2 (Proteintech, 10412-1-AP), rabbit anti-MLH1 (Cell Signaling Technology, 3515T), rabbit anti-H2A.Z (Abcam, ab4174).

### *Znhit1* in situ hybridization

Testis sections (5 μM) at the indicated times were prepared for *Znhit1* in situ hybridization with the RNAscope kit (Advanced Cell Diagnostics, 323100) according to the manufacturer’s instructions.

### Quantitative reverse transcription PCR (RT-qPCR)

Total RNA was isolated using the RNeasy Mini-plus Kit (Qiagen, 74134) and then reverse-transcribed into complementary DNA (cDNA) with the GoScript Reverse Transcription System (Promega, A5003). cDNA was used as the template for the quantitative PCR assay using 2×SYBR Green qPCR Master Mix (Bimake, B21202). Quantitative PCR primers are listed in Table S7.

### RNA-seq library generation and sequencing

Total RNA from fresh testes was isolated and mRNA was purified with magnetic beads (Vazyme, N401-01). Then, mRNA was fragmented and processed to generate RNA-seq libraries (Vazyme, NR605-01). Over 40 million reads were obtained per sample using the Illumina NovaSeq platform for 2 independent biological replicates.

### Single-cell RNA sequencing (scRNA-seq)

P16 and P35 control and *Znhit1*-sKO testes were digested by collagenase IV and trypsin at 37D for 10 min to obtain testicular cell suspensions. scRNA-seq libraries were constructed using a 10x Genomics kit and sequenced on the Illumina platform.

### ChIP-seq library generation and sequencing

ChIP-seq library generation was performed as previously described. Briefly, P14 control and *Znhit1*-sKO testes were digested by collagenase IV and trypsin at 37D for 10 min, crosslinked with 1% formaldehyde for 10 min, and quenched with glycine. The cells were lysed and sheared with a Bioruptor Plus machine for 20 min. Then, 2 μg of anti-H2A.Z antibody (Abcam, ab4174) was added to the sonicated chromatin and incubated overnight at 4D. The following day, 20 μL of protein G beads were added and incubated for 2 h at 4D. The beads were washed and reverse-crosslinked for 4 h at 65D. DNA was extracted using phenol-chloroform and subjected to library construction using the VAHTS Universal DNA Library Prep Kit for Illumina (Vazyme, ND607-01) according to the manufacturer’s instructions. Over 30 million reads were obtained per sample using the Illumina NovaSeq platform for 2 independent biological replicates.

### KAS-seq library and ATAC-seq library generation and sequencing

Testes were digested by collagenase IV and trypsin at 37D for 10 min. Pachytene cells were sorted using fluorescence-activated cell sorting (FACS) as previously described (Long et al., 2017). For KAS-seq, pachytene cells were labeled with N3-kethoxal, and DNA was isolated using the DNA Clean and Concentrator kit (Zymo, D4013) and subjected to library construction using Q5 high-fidelity DNA polymerase (New England Biolabs, M0544S). Genomic DNA was fragmented using Tn5 transposase and subjected to library construction. For ATAC-seq, pachytene nuclei were isolated and fragmented with Tn5 transposase. DNA was isolated using the DNA Clean and Concentrator kit (Zymo, D4013) and subjected to library construction using Q5 high-fidelity DNA polymerase (New England Biolabs, M0544S). For high-throughput sequencing, over 30 million reads were obtained per sample using the Illumina NovaSeq platform for 2 independent biological replicates.

### RNA-seq data analysis

We performed RNA-seq analysis as described previously (Pertea et al., 2016). Briefly, after removing adapters using Cutadapt (v2.5) (Kechin et al., 2017), paired-end reads were aligned to the annotated mouse transcripts (mm10 Gencode vM23 release) using Hisat2 (v2.2.1) (Kim et al., 2015; Kim et al., 2019). Gene expression levels were calculated using StringTie (v2.2.1) (Pertea et al., 2015). Read counts were calculated using a Python script (prepDE.py3) provided by the StringTie development team. Differentially expressed genes were identified using the R package DESeq2 (v1.38.3) (Love et al., 2014). Genes with read counts > 50 in at least one sample were kept for further analysis. A given gene was considered to significantly changed if the adjusted P value (padj) was < 0.05, the P value was < 0.01, and the fold-change was≥ 2. Gene Set Enrichment Analysis (GSEA) was carried out using GSEA software (Subramanian et al., 2005). GO analysis was performed using clusterProfiler (v4.6.2) (Wu et al., 2021).

### scRNA-seq data analysis

FASTQ files were run through CellRanger (v7.1.0) software with default parameters for de-multiplexing, aligning reads with STAR software to mm10, and counting unique molecular identifiers (UMIs). As input files of the Seurat R package (v4.4.0), the filtered gene expression matrices were then used for downstream analyses (Butler et al., 2018). Low-quality cells were filtered (expressing < 500 or >6,000 unique gene counts and >15% mitochondrial reads). Principal component analysis was performed on SCT-transformed data using 3,000 variable genes. The top 50 principal components were used for clustering and visualized using the UMAP algorithm in the Seurat R package. The "FindAllMarkers" function of the Seurat R package was used to calculate cluster-specific genes. Marker genes for each cluster are shown in Table S8.

The "FindMarkers" function of the Seurat R package was used to identify differentially expressed genes (DEGs) for spermatocytes (preleptotene, leptotene, zygotene, and pachytene). Only those with |’avg_logFC’| > 0.25 and ‘p_val_adj’ < 0.05 were considered as DEGs. For the transcriptional regulatory network analysis, the raw count matrix was used by the pyscenic (v0.12.1) workflow using default parameters (Aibar et al., 2017). Then, the output loom file was opened by the SCENIC R package (v1.3.1), and AUC values of regulons were extracted for visualization of downstream transcription factor activity.

### ChIP-seq and KAS-seq data analysis

For analyzing ChIP-seq and KAS-seq data, we used the ENCODE ChIP-seq pipeline (v2.2.1, https://github.com/ENCODE-DCC/chip-seq-pipeline2). Raw reads were cleaned by Cutadapt (v2.5) (Kechin et al., 2017). After removing adapters, clean reads were aligned against the mouse mm10 genome using bwa (v0.7.17) (Li and Durbin, 2009). Then, sam files were converted to bam files using samtools (v1.9) (Li et al., 2009). PCR duplicates were removed using Picard (v2.20.7) (https://broadinstitute.github.io/picard/). Histone ChIP-seq peaks (such as H2A.Z and histone marks) and KAS-seq peaks were called using MACS2 (v2.2.4) (Zhang et al., 2008), while transcription factor ChIP-seq peaks (such as A-MYB) were called using the R packages spp (v1.15.5) (https://github.com/hms-dbmi/spp). The conservative narrow peaks were used for downstream analysis.

For visualization, bam files were converted to bigWig files using deepTools (v3.3.1) (Ramirez et al., 2014), and bigWig files were used to calculate tag density under 50 bp resolution. In addition, CPM was used to normalize the number of reads per bin. We used the R package DiffBind (v3.8.4) (http://bioconductor.org/packages/release/bioc/html/DiffBind.html) for differential peak analysis and the R package rGREAT (v2.1.11) (Gu and Hubschmann, 2023) for peak annotation. We used ComputeMatrix and plotHeatmap of deepTools to generate count matrices and heatmaps, respectively. H2A.Z enrichment within chromatin states was annotated using the ChromHMM model (Ernst and Kellis, 2012; Spruce et al., 2020). TF motif enrichment was calculated using HOMER (v4.11) (Heinz et al., 2010). We used deepTools to draw Pearson’s correlation scatter plots for the correlation between replicates of each experiment.

For promoter and enhancer type analysis, H3K27ac ChIP-seq data in wild-type pachytene spermatocytes were downloaded from the published data (GSE107398) (Adams et al., 2018); ChIP-seq data for H3K4me1, H3K4me3, and H3K27me3 in wild-type pachytene spermatocytes were downloaded from the published data (GSE132446) (Chen et al., 2020).

### ATAC-seq data analysis

ATAC-seq data were analyzed using a standardized ENCODE ATAC-seq pipeline (v2.2.2, https://github.com/ENCODE-DCC/atac-seq-pipeline). The mouse reference genome (mm10) was used in the pipeline. Trimming, mapping, and duplicate-removing were performed as the ENCODE pipeline suggested using Cutadapt (v2.5) (Kechin et al., 2017), Bowtie2 (v2.3.4.3) (Langmead et al., 2009), and Picard (v2.20.7) (https://broadinstitute.github.io/picard/) respectively. MACS2 (v2.2.4) (Zhang et al., 2008) was used for peak calling.

### Statistics and reproducibility

Statistical analyses were performed using R and Prism 9. Data are presented as means ± s.d. unless otherwise indicated. The two-tailed, unpaired, Student’s t-test, Mann–Whitney test, or one-way or two-way ANOVA were performed to analyze statistical significance.

## Supplementary Materials

This manuscript file includes Figs. S1 to S9 and Tables S1 and S8.

## Acknowledgments

We thank Chuan He for N3-kethoxal and Christopher L. Baker for meiotic chromatin state data.

## Funding

This work was supported by grants from the National Key Research and Development Program of China (2022YFA0806200, 2018YFC1003500, 2018YFA0800100, 2021YFC2501800), the National Natural Science Foundation of China (32192400, 81971443, 32300702, 32350710191), the Science and Technology Major Project of Inner Mongolia Autonomous Region of China to the State Key Laboratory of Reproductive Regulation and Breeding of Grassland Livestock (2020ZD0008). Shenfei Sun was supported by the fellowship of China Postdoctoral Science Foundation (2022M720797) and the Postdoctoral Fellowship Program (Grade B) of China Postdoctoral Science Foundation (GZB20230161).

## Author Contributions

S.S. and X.L. conceived and designed the study; S.S. performed most of the experiments with the help of Y.J., Q.Z., H.P., F.H., X.Z., Y.G., X.Y., K.G., W.W., H.L., Z.S., Y.S., X.T., M.Y., and R.L.; Y.J. and N.J. performed the bioinformatics analysis; X.L. supervised the work and acquired the funding support; and S.S. and X.L. wrote the manuscript, with contributions from all authors.

## Competing interests

The authors declare that they have no competing interests.

## Data and materials availability

The NGS data generated in this study were deposited in the NCBI SRA database under accession numbers SRP467214 (RNA-seq and ChIP-seq data) and SRP467448 (ATAC-seq and KAS-seq data).

## Supplementary Materials

**Figure S1.**
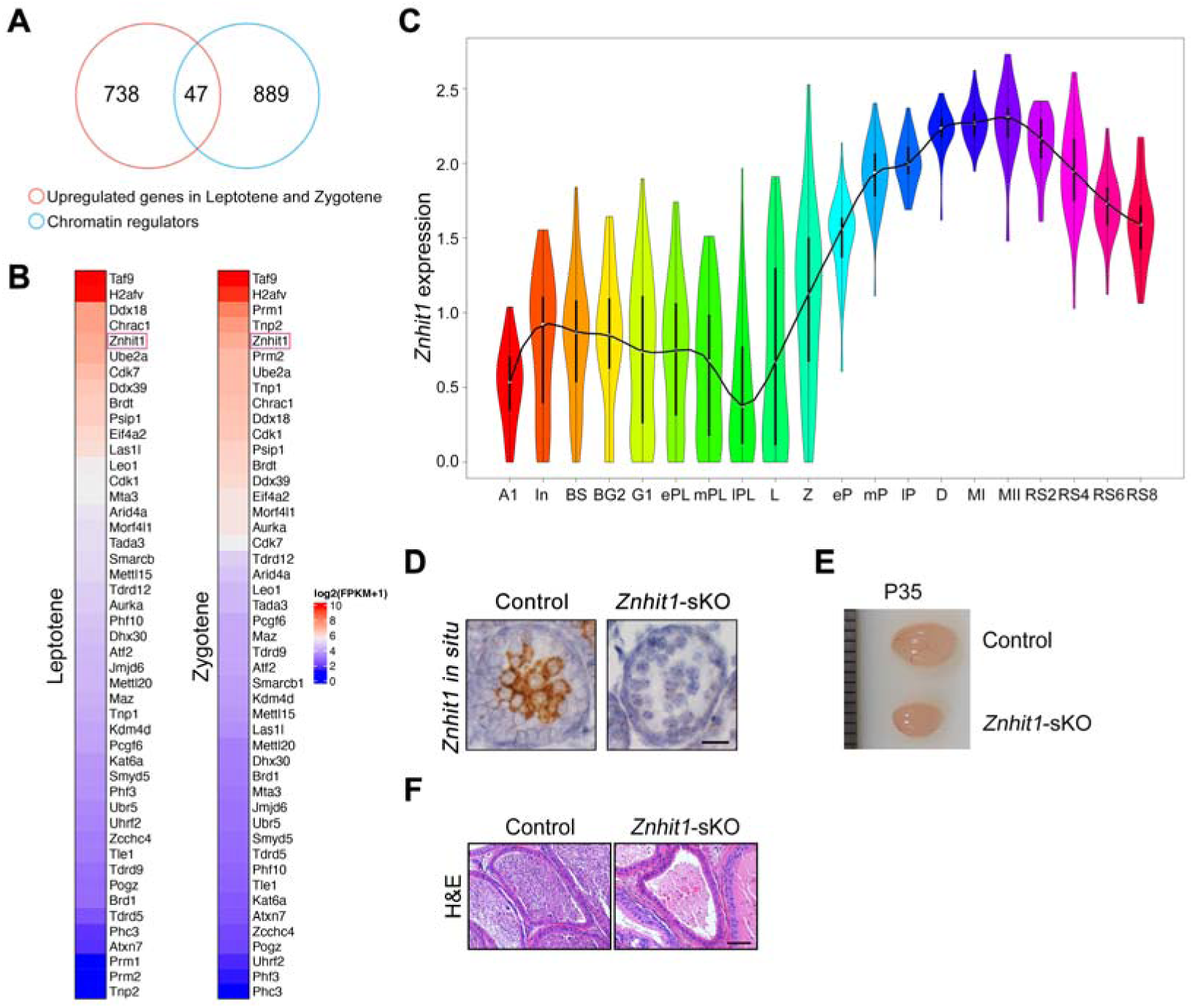
*Znhit1* expression during meiotic progression and spermatocyte-conditional deletion of *Znhit1* with Stra8-cre. (**A** and **B**) Venn diagram and heatmaps showing chromatin regulators highly expressed during leptotene and zygotene spermatocytes. (**C**) Violin plot showing *Znhit1* transcription level during spermatogenesis. Gene expression profiles derived from published scRNA-seq data (Chen et al, 2018). (**D**) *Znhit1* in situ hybridization in P14 testis sections. Scale bar, 20 μm. (**E**) Testis size comparison between control and *Znhit1*-sKO mice at P35. (**F**) Hematoxylin and eosin (H&E) - stained testis sections from control and *Znhit1*-sKO mice at P60. Scale bar, 50 μm.

**Figure S2.**
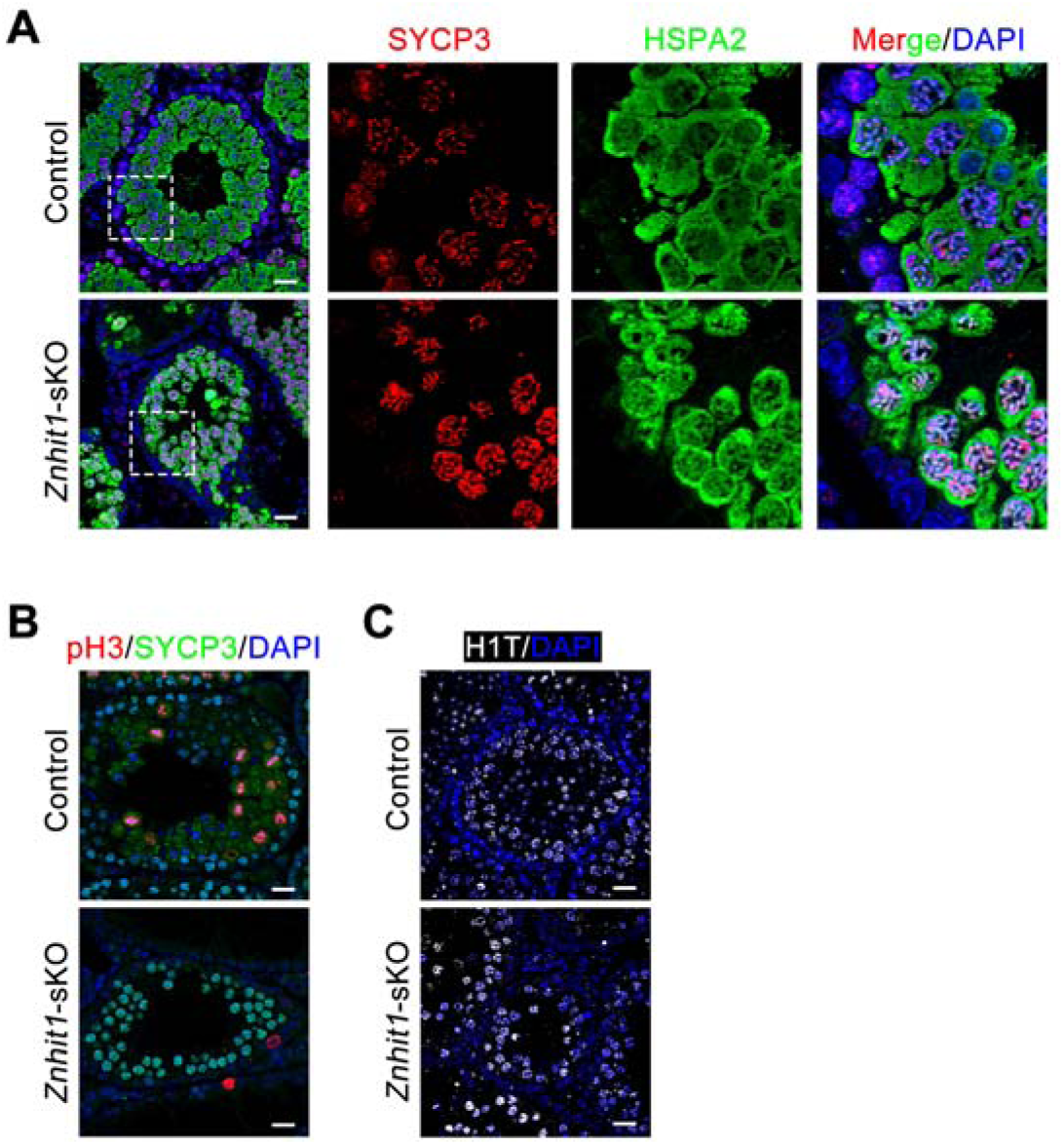
Spermatocyte development in *Znhit1*-sKO testes. (**A**) Immunostaining of SYCP3 and HSPA2 in testis sections of control or *Znhit1*-sKO mice at P35. Scale bar, 20 μm. (**B**) Immunostaining of SYCP3 and pH3 in testis sections of control or *Znhit1*-sKO mice at P35. Scale bar, 20 μm. (**C**) Immunostaining of H1T in testis sections from control and *Znhit1*-sKO mice at P35. Scale bar, 20 μm.

**Figure S3.**
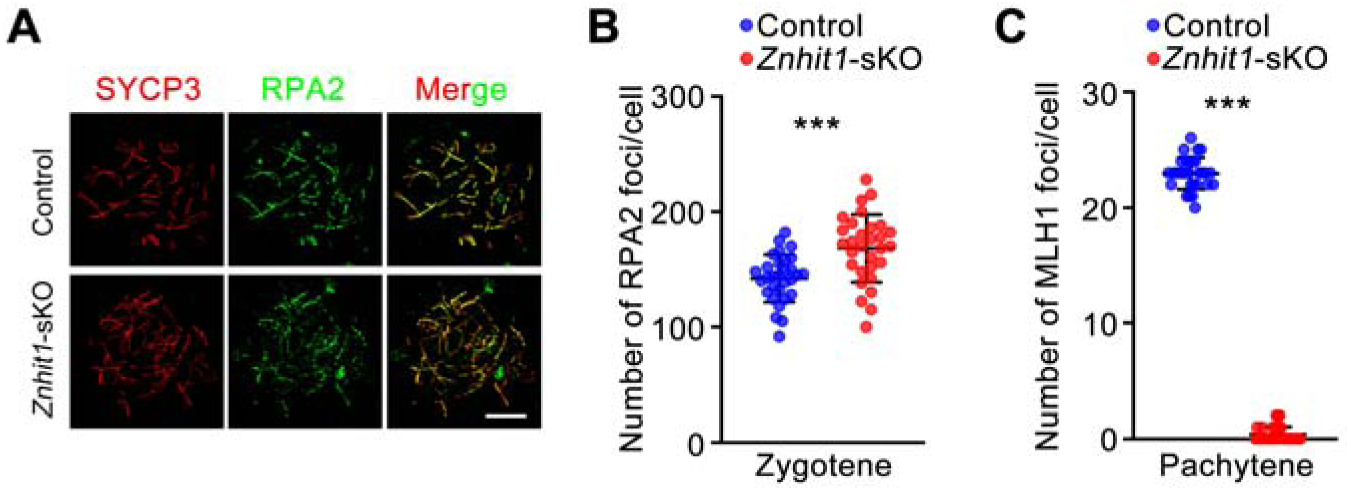
ZNHIT1 is required for meiotic recombination. (**A** and **B**) Immunostaining of SYCP3 and RPA2 in spermatocyte chromosome spreads of control or *Znhit1*-sKO mice. Quantitative data are shown in (**B**). Scale bar, 10 μm. (**C**) Plots showing quantitative data of MLH1 foci of control or *Znhit1*-sKO mice. Data are presented as the mean ± s.d. *** *p* < 0.001

**Figure S4.**
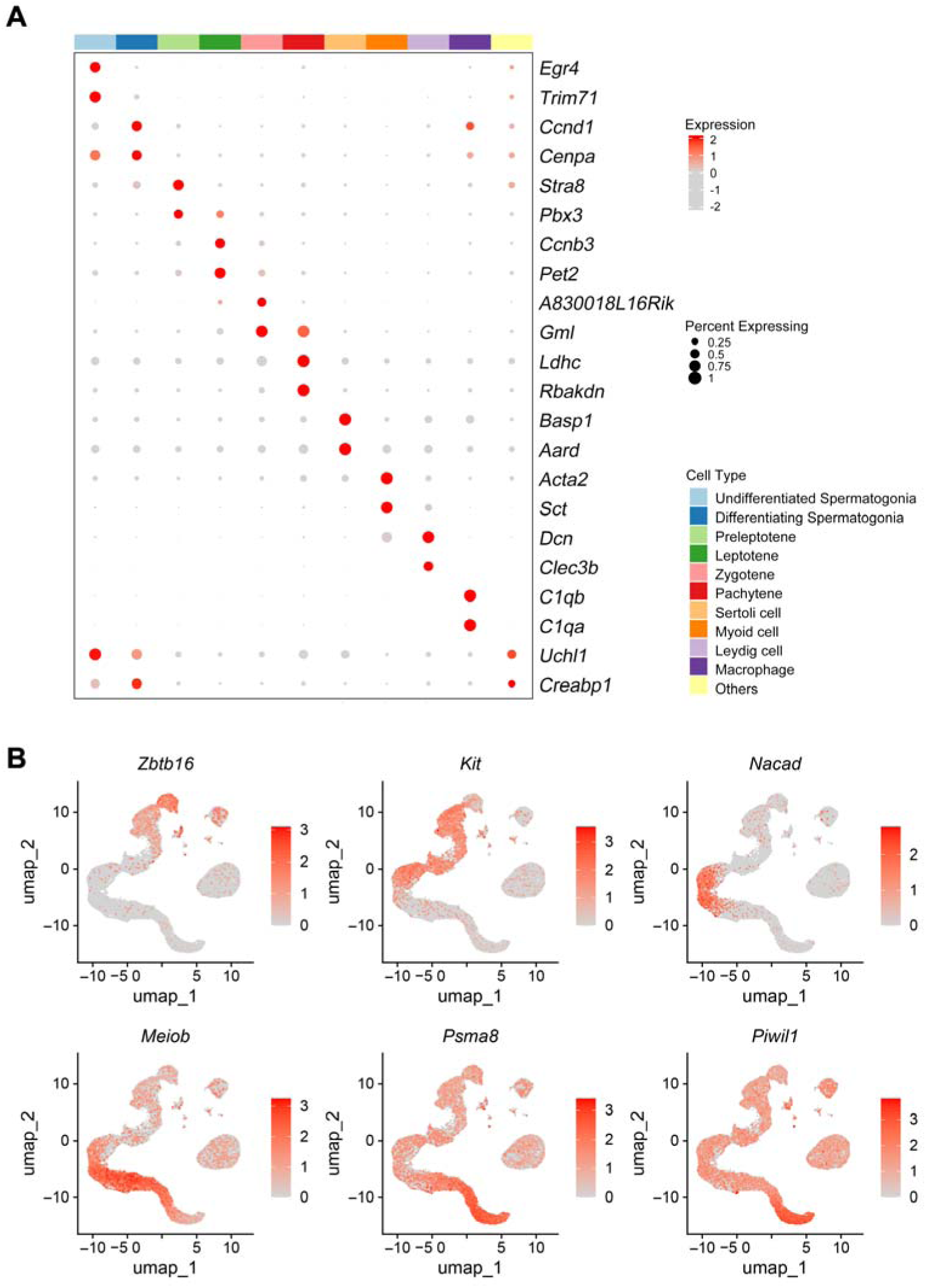
Single-cell RNA-seq of testicular cells. (**A**) Dot plot showing the relative expression and the percentage of cells expressing selected markers across scRNA-seq clusters. (**B**) UMAP plots showing the normalized expression of the indicated genes in scRNA-seq data.

**Figure S5.**
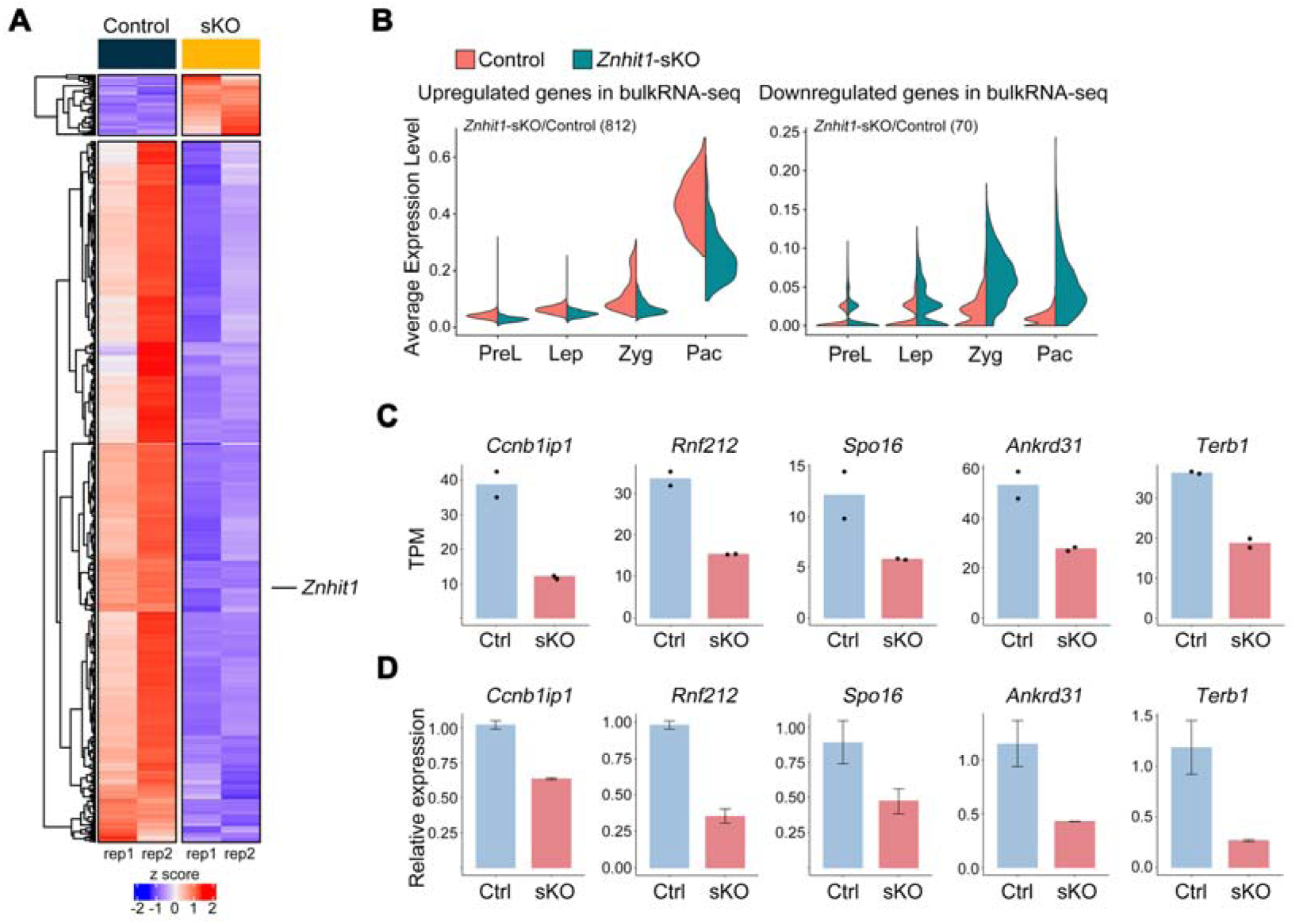
Transcriptomic analysis of testicular cells. (**A**) Heatmap showing differentially expressed genes (DEGs) in P14 testicular cells between control and *Znhit1*-sKO mice. (**B**) Violin plots showing the average expression level of upregulated or downregulated genes between P14 control and *Znhit*1-sKO testicular cells. PreL, preleptotene; Lep, leptotene; Zyg, zygotene; Pac, pachytene. (**C** and **D**) Bar charts showing HR-related gene expression in P14 control and *Znhit*1-sKO testes. Data from bulk RNA-seq are shown in (**C**). Validated data from RT-qPCR are shown in (**D**). Ctrl, control; sKO, *Znhit1*-sKO.

**Figure S6.**
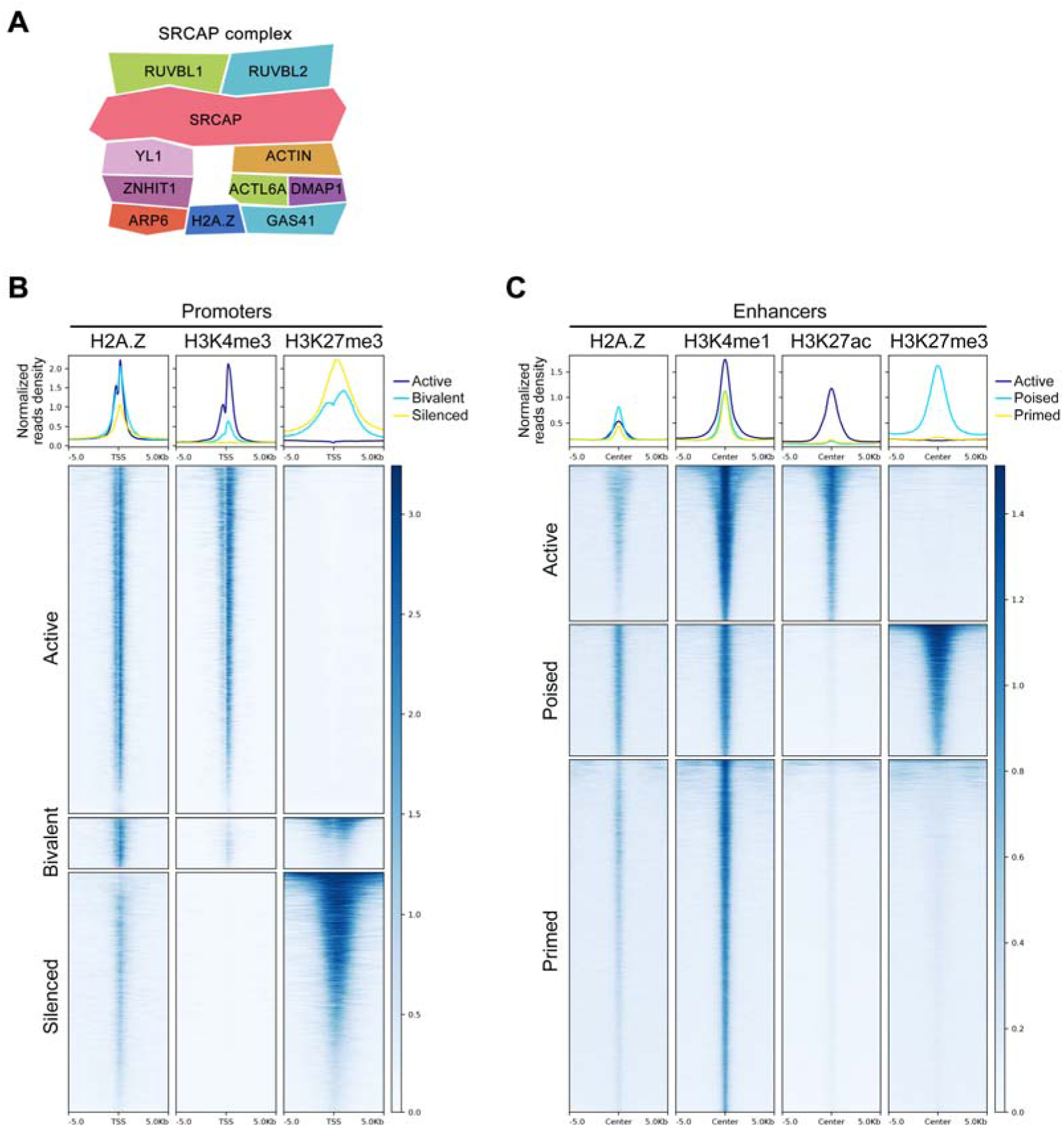
H2A.Z is enriched at meiotic gene promoters and enhancers. (**A**) Schematic representation of the components of the SRCAP chromatin remodeling complex. (**B**) Heatmaps showing H2A.Z ChIP-seq signals surrounding different types of promoters. (**C**) Heatmaps showing H2A.Z ChIP-seq signals surrounding different types of enhancers.

**Figure S7.**
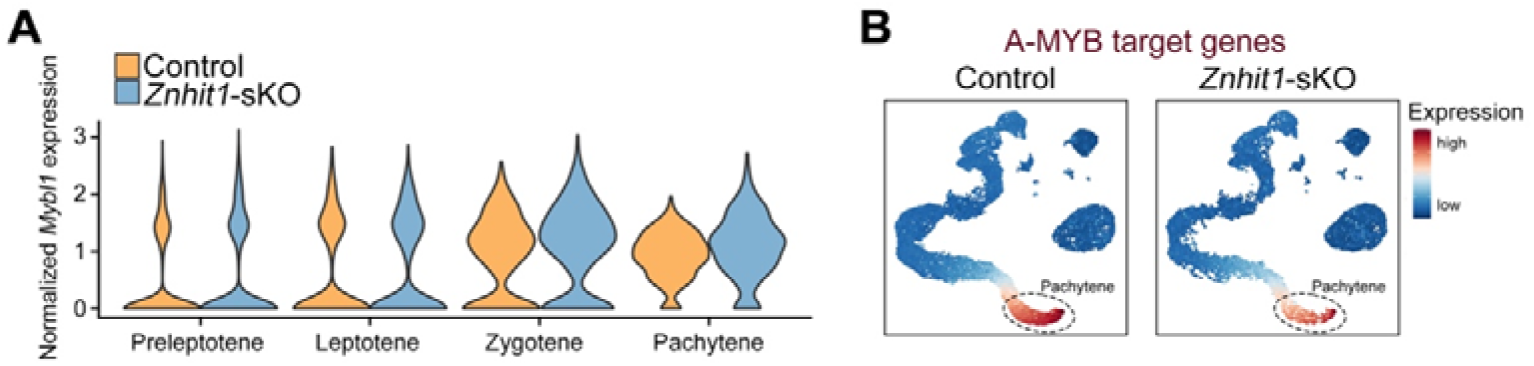
*Znhit1* deletion downregulates the expression of A-MYB target genes. (**A**) Violin plot showing the normalized expression of *Mybl1* in control and *Znhit1*-sKO spermatocytes at consecutive stages. (**B**) Cluster annotation on the basis of the expression of A-MYB target genes in control or *Znhit1*-sKO testicular cells.

**Figure S8.**
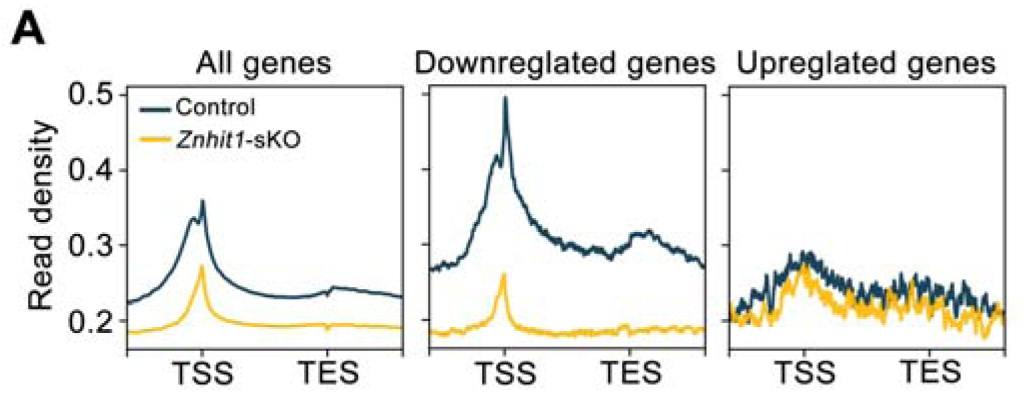
KAS-seq analysis. (**A**) Mean plots of KAS-seq showing signal changes within indicated genes.

**Figure S9.**
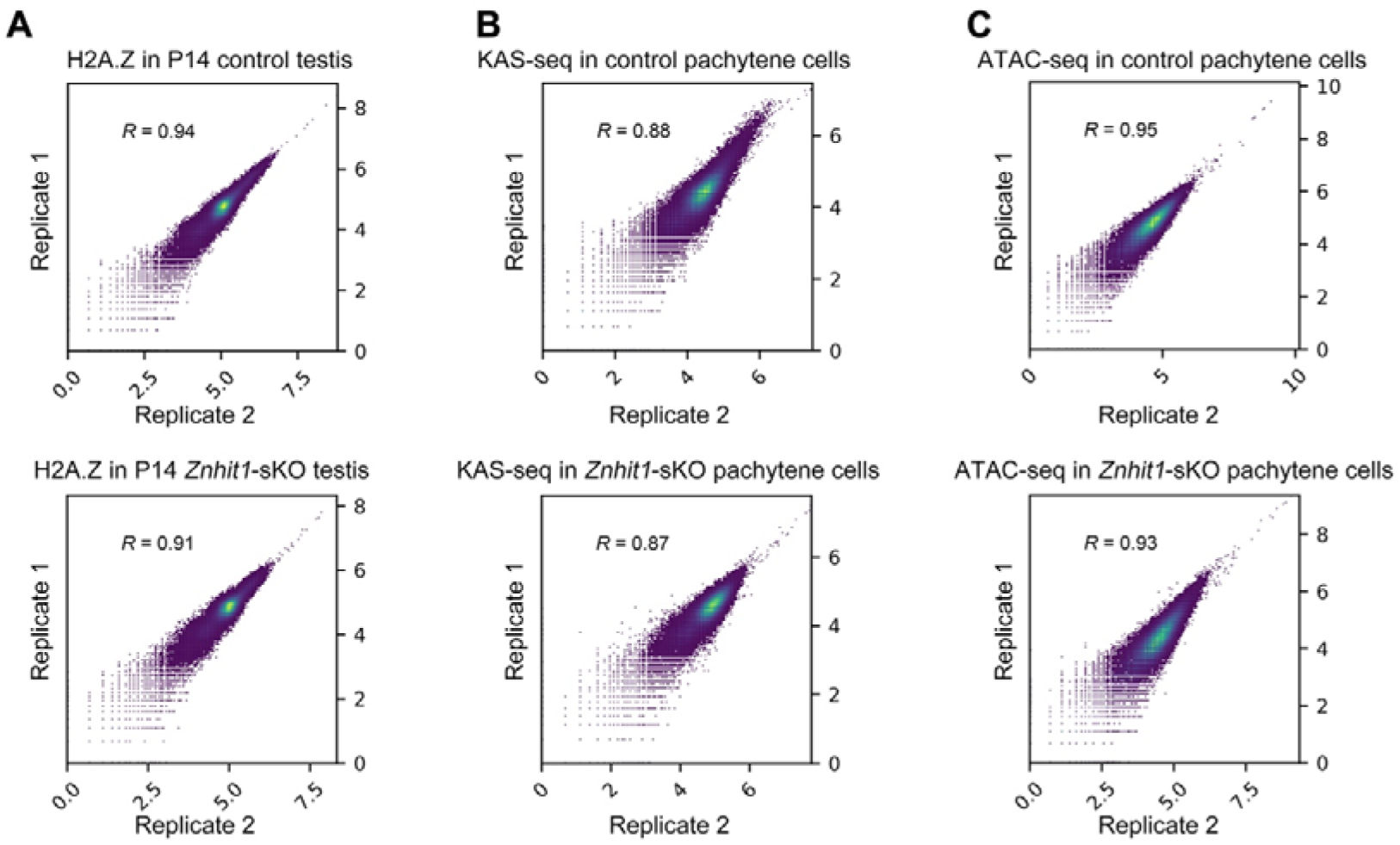
Reproducibility of replicates in this study. (**A**) Scatter plots showing H2A.Z enrichments in P14 control or *Znhit1*-sKO testes for two independent H2A.Z ChIP-seq replicates. Pearson correlation coefficients are indicated. (**B**) Scatter plots showing KAS-seq signals in P14 control or *Znhit1*-sKO pachytene cells for two independent H2A.Z ChIP-seq replicates. Pearson correlation coefficients are indicated. (**C**) Scatter plots showing ATAC-seq signals in P14 control or *Znhit1*-sKO pachytene cells for two independent H2A.Z ChIP-seq replicates. Pearson correlation coefficients are indicated.

## Notes

### Competing Interest Statement

The authors have declared no competing interest.

### Summary of Updates

Title, abstract, and manuscript have been revised following the comments of reviewers.

